# Higher glucose levels buffer against everyday stress load

**DOI:** 10.64898/2025.12.08.692911

**Authors:** Madeleine Kördel, Anne Kühnel, Kristin Kaduk, Alica Guzman, Melanie Henes, Melina Grahlow, Birgit Derntl, Nils B. Kroemer

## Abstract

Adaptive stress responses are dependent on the availability of energy and the body’s effectiveness in metabolizing glucose as fuel. However, it is not well understood if glucose levels contribute to the functional regulation of daily stress in healthy individuals. Here, we used 4,307 ratings of stress during a four-week period of ecological momentary assessment (EMA) with concurrent recordings from continuous glucose monitoring (CGM) in 91 participants (47 women). In accordance with the hypothesized metabolic link, we found that higher glucose levels were associated with lower self-reported stress (*b* = −.038, *p* = .035, controlling for mood). Crucially, dynamic changes in glucose levels also precipitated subsequent increases in stress levels (0-2h before EMA: *p* = .013; incremental effect 10-12h before: *p* < .001), with lower glucose levels associated with subsequent stress increases and elevated glucose with stress relief, highlighting the potential of monitoring physiological biosensor data to improve adaptive stress regulation. We conclude that metabolic signals, such as glucose, play an important role in channeling stress responsiveness in healthy adults, pointing to a pivotal avenue for improved interventions to avoid excessive stress experiences.

## Introduction

Stress is an energetically demanding state that triggers neuroendocrine responses scaling with glucose availability (Kördel et al., 2025). In response to physical or psychological challenges, the acute stress response mobilizes energy (Chrousos & Gold, 1992; Sapolsky et al., 2000) via activation of the hypothalamic-pituitary-adrenal (HPA) axis and the sympathetic nervous system (Goel et al., 2014; Lupien et al., 2005), leading to the release of cortisol (Marin et al., 2011). To ensure energy availability for the brain and body, cortisol raises blood glucose levels by promoting gluconeogenesis, reducing glucose uptake by tissues, and inhibiting insulin secretion (Russell & Lightman, 2019; Sapolsky et al., 2000). While acute stress-induced hyperglycemia supports short-term adaptation, repeated or prolonged activation of these systems increases allostatic load, contributing to adverse outcomes, including metabolic and mental disorders (Marin et al., 2011; McEwen, 1993; Russell & Lightman, 2019) and leading to additional energetic costs for the organism (Bobba-Alves et al., 2022). Although the effects of stress on glucose metabolism are well established, how glucose availability modulates stress responses is less well understood.

Emerging evidence shows that glucose levels affect acute stress responses, suggesting that the relationship between stress and glucose is bidirectional. Most research has focused on physiological stress markers, showing that glucose availability shapes HPA axis responses. Experimental studies in healthy participants demonstrate that cortisol reactivity is increased with a prior glucose but not fat or protein load (Bentele et al., 2021; Gonzalez-Bono et al., 2002; Kirschbaum et al., 1997; Kördel et al., 2025; von Dawans et al., 2021). In addition, central insulin modulates HPA axis activity, with intranasal insulin dampening cortisol responses to psychosocial stress (Bohringer et al., 2008). Further acute and chronic glucose exposure may affect stress physiology in opposing ways, as habitual sugar intake is associated with attenuated cortisol response to stress (Di Polito et al., 2023). This aligns with the “comfort food” theory, which proposes that repeated intake of palatable foods may downregulate HPA axis responsiveness over time (Tomiyama et al., 2011). Likewise, a higher BMI is associated with altered HPA axis functioning (Kühnel et al., 2023), including reduced cortisol response to stress and lower baseline cortisol levels in individuals with obesity compared to those with normal weight (Herhaus et al., 2025; Herhaus & Petrowski, 2018).

In contrast, the evidence for the effects of glucose on subjective stress and mood is more mixed. In daily life, individuals with type 2 diabetes report greater emotional reactivity to stress when fasting blood glucose levels are elevated (Rook et al., 2016). In experimental settings, changes in subjective stress have not been observed after a glucose load (von Dawans et al., 2021), although most studies did not test whether glucose administration also increased the subjective stress experience (Bentele et al., 2021; Gonzalez-Bono et al., 2002; Rüttgens & Wolf, 2022; Zänkert et al., 2020). Nonetheless, earlier experimental work has suggested that ingestion of a glucose drink and subsequent higher blood glucose levels can buffer negative mood and tension for a short time in healthy individuals (Benton & Owens, 1993). To summarize, converging evidence suggests that glucose availability influences physiological stress reactivity in lab-based settings, but the association with subjective stress experiences in everyday life has not been examined in metabolically healthy adults.

Although controlled laboratory manipulations provide valuable mechanistic insights, they do not address whether energy metabolism modulates naturalistic stress experiences. Such approaches have become feasible only recently with the availability of continuous glucose monitoring (CGM). The integration of CGM with repeated ecological momentary assessments (EMA) enables measuring glucose variations independently of food intake or exercise (Holzer et al., 2022; Kaduk et al., 2025; Schrems et al., 2025), while tracking subjective experiences as they naturally unfold. So far, studies using CGM have been conducted primarily in populations with diabetes, and they report mixed associations between glucose, negative affect, and quality of life (Muijs et al., 2021; Penckofer et al., 2012), without distinguishing stress experiences. Some of the studies combined CGM with EMA (De Wit et al., 2023; Muijs et al., 2021; Penckofer et al., 2012; Schrems et al., 2025), typically covering only short monitoring periods of a few days up to two weeks. Recent work has extended this approach to non-diabetic populations to explore the dynamic relationships between glucose fluctuations and affective states, demonstrating the methodological feasibility of this approach outside of clinical settings (Kaduk et al., 2025; Rethorst et al., 2023; Zink et al., 2020). However, these studies have not addressed stress or stress-related parameters.

Taken together, while the effects of stress on glucose metabolism are well established, less is known about the reverse association and the role of glucose availability in shaping subjective stress experiences in everyday life of healthy individuals. To address this gap, we combined four weeks of CGM with EMA, capturing self-reports of subjective stress in naturalistic settings (Shiffman et al., 2008) in 90 mentally and metabolically healthy participants. We tested the hypotheses that elevated glucose levels would be associated with lower perceived stress under naturalistic conditions, and that differences in glucose levels would precede changes in self-reported stress. Consistent with our predictions, higher glucose availability was linked to reduced daily stress load, whereas short-term glucose dynamics precipitated increases or decreases in stress, highlighting a pivotal interaction between metabolic and psychological processes.

## Methods

We preregistered the study at the Open Science Framework (https://osf.io/gpr52) and the results are part of a larger study examining the association between metabolic states and reward learning. The specific analyses were not preregistered, and the study design was previously reported in Kaduk et al. (2025).

### Participants

For the current analyses, the sample included 91 participants (47 women, *M*_age_: 24.24 ± 3.56 years, range: 18–34, *M*_BMI_: 24.74 ± 4.09 kg/m^2^, range: 18-36.7). Mean age and BMI did not differ between female and male participants. In total, 97 participants were enrolled, of whom 6 were excluded due to missing CGM data (e.g., detached sensors or loss of transmission). No dietary restrictions were enforced. Participants were included in the final sample if they completed ≥20 EMA runs with concurrent CGM data (Kaduk et al., 2025). According to phone screenings, all participants were healthy (except for BMI) and reported no history of neurological, neurosurgical, or cardiological disease or treatment. They received fixed compensation of 160€ or partial course credits and performance-dependent wins from the tasks and EMA. The sample size was selected to provide at least a power of 1-*β* = .94 to study small-to-moderately sized within-subject effects (*d*_z_∼.40), leading to a lower-bound estimate of 80 participants after quality control. All participants provided their written informed consent before the experiment. The ethics committee of the Faculty of Medicine at the University of Tübingen approved the experiment, and all procedures were carried out in accordance with the Declaration of Helsinki.

### Experimental procedure

Participants completed five weekly laboratory sessions over one month (Fig. 1), followed by two MR sessions (not reported here). In the first session (S1), participants arrived after an overnight fast (>12h) and completed three tasks (effort allocation (Neuser et al., 2020), go/nogo reinforcement learning (Kühnel, Teckentrup, et al., 2020), reward ratings (Müller et al., 2022), and state questions) after a blood draw to determine fasting levels of insulin, glucose, and triglycerides. During the first session, participants were equipped with a CGM sensor (FreeStyle Libre 3 sensors, Abbott GmbH, Abbott Diabetes Care, Wiesbaden, Germany) and received a cell phone to track their food intake, synchronize the CGM data, and play a reward game (Influenca; see Neuser et al., 2023). EMA questions covered stress, mood (happy and sad), and metabolic (hungry and sated) states and were embedded in the completion of the gamified Influenca task (see Kaduk et al., 2025). For the stress and state ratings, participants indicated their response on a visual analog scale (VAS), ranging from 0, representing "not at all", to 100, representing "very much”. Participants were asked to play twice a day over four weeks with a minimum latency of 2 h (to sample different states). After two weeks, during the third lab session, the first glucose sensor was replaced. Faulty CGM sensors were replaced if necessary. At the MR sessions, we again drew blood after an overnight fast to measure fasting glucose, insulin, and triglyceride levels.

**Figure 1.**
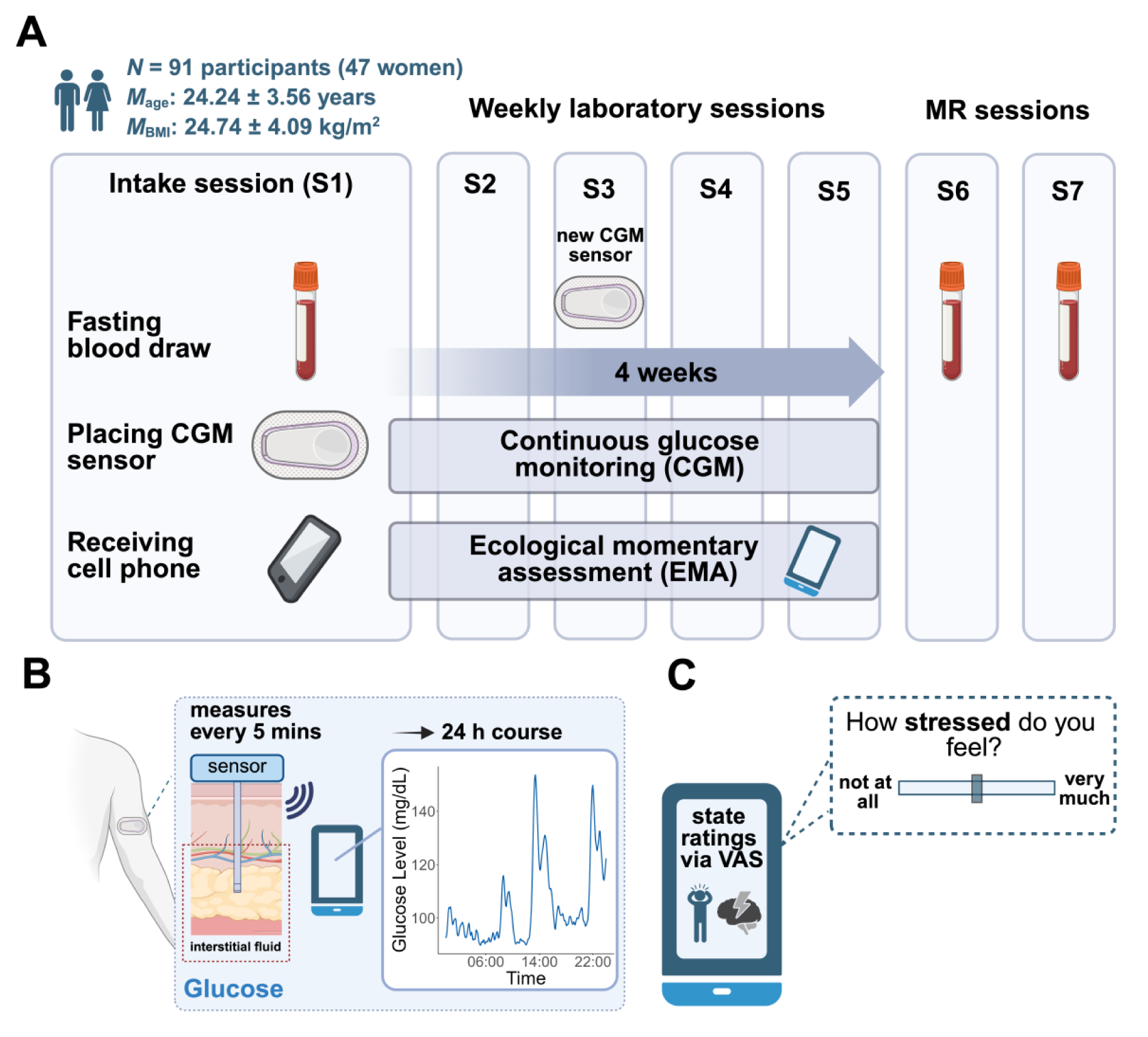
Summary of the study. **A:** The schematic shows the experimental procedure, illustrating five weekly lab visits followed by 2 MR visits labeled from S1 to S7. After the first session (S1), participants underwent four weeks of continuous glucose monitoring (CGM) combined with ecological momentary assessment (EMA). **B:** The CGM system uses a small flex sensor placed at the posterior upper arm that is inserted in the subcutaneous fatty tissue. The glucose sensor, positioned beneath the skin surface layers (epidermis and dermis), measures glucose levels in the interstitial fluid within the subcutaneous adipose tissue. The data is wirelessly sent via near field communication (NFC) to the FreeStyle Libre 3 app on the participant’s cell phone. The accuracy of the sensor concerning capillary glucose has been validated (Alva et al., 2022). **C:** Participants rate their metabolic and stress states twice daily using a visual analog scale (VAS), ranging from 0 (not at all) to 100 (very). Created with BioRender.com (adapted from Kaduk et al., 2025).

### Data analysis

#### Preprocessing of CGM data

We extracted raw sensor glucose values and preprocessed them according to best-practice recommendations (Visser & Gillard, 2024) as previously described (Kaduk et al., 2025). We segmented the data into 20-minute intervals and cleaned it using MATLAB’s ‘msbackadj’ function from the Bioinformatics toolbox. Briefly, for each participant, we identified a baseline glucose period with the lowest glucose variability using the raw CGM during the overnight fasting window (midnight to 8 am). After baseline correction, CGM data was smoothed with a Gaussian kernel and a window length of 5 recorded values. We then partitioned the adjusted data into standardized five-minute intervals. As the physiological delay of glucose transport from the vascular to the interstitial space is about 5 to 6 min in healthy adults (Basu et al., 2013), we extracted the CGM value, which was recorded 5-10 min after the corresponding EMA as the concurrent glucose level. The corresponding MATLAB code is available on GitHub (https://github.com/neuromadlab/Glucose_Stress).

#### Insulin resistance derived from blood samples

As measures of insulin sensitivity, we calculated the homeostasis model assessment of insulin resistance (HOMA-IR; higher HOMA-IR reflects lower insulin sensitivity) using the glucose and insulin levels measured from fasting blood samples (Matthews et al., 1985). As an additional measure of insulin resistance, we calculated the triglyceride-glucose (TyG) index, based on fasting triglyceride and glucose concentrations (Simental-Mendía et al., 2008). Higher TyG values indicate lower insulin sensitivity. Most participants had three fasting blood samples (baseline session and both MRI sessions) to compute the median HOMA-IR, though one participant had only two samples, and nine participants had only one. Since the distribution of the HOMA-IR and glucose levels was skewed, we ln-transformed them for parametric analyses (Emoto et al., 1999; Kroemer et al., 2013). We also residualized HOMA-IR values by adjusting for BMI, sex, and age to evaluate the unique association with insulin sensitivity.

#### Statistical analyses and software

To determine how strongly stress ratings were associated with mood, we predicted stress with grand mean-centered mood state as the predictor. To evaluate whether this association differed for positive and negative mood items, we predicted stress with grand mean-centered happy and sad ratings as predictors. Mood state was computed as a composite of positive and negative mood ratings, with the negative mood item (sad) subtracted from the positive mood item (happy) (Kaduk et al., 2025). Next, to quantify short-term fluctuations in stress and mood, we computed temporal derivatives of stress and mood by calculating differences between consecutive ratings within each participant. Then, to examine the relationship between changes in stress and mood, we predicted changes in stress with *z*-standardized individual averages of mood state and moment-to-moment changes in mood as predictors. To evaluate whether within-person fluctuations in mood predicted changes in stress, we predicted changes in stress with grand mean-centered mood state and changes in mood as predictors. All models included BMI and age as grand mean-centered covariates and sex as an effect-coded covariate (male = 0.5, female = −0.5).

To assess whether concurrent glucose levels explain additional variance in stress beyond the current mood state, we predicted stress with group-centered (i.e., per ID) glucose levels, individual averages of glucose, and grand mean-centered happy and sad as predictors. In addition, the model included interaction terms between group-centered glucose and both mood items. To test whether changes in glucose preceding EMA were associated with stress changes, we predicted estimated glucose levels using six 2-hour time intervals (bins) relative to the EMA stress rating (from 12 to 2 hours before the rating, dummy-coded with reference category 0-2h before the EMA), *z*-standardized changes in stress, and their interaction. To account for potential circadian effects, we added grand mean-centered hour of day as a covariate. Additionally, to examine whether stress levels varied by time of day, we predicted the hour of EMA stress ratings with *z*-standardized change in stress as a predictor.

To examine the role of BMI in stress dynamics, we classified participants into three categories: normal weight (BMI < 25 kg/m²), overweight (BMI 25-29.9 kg/m²), and obese (BMI ≥ 30 kg/m²) (WHO, 1995). This resulted in 54 participants with normal weight, 27 with overweight, and 10 with obesity.

We preprocessed CGM data with MATLAB vR2022b. All statistical analyses were conducted with R (v4.3.1, R Core Team, 2023) and linear mixed-effects models were estimated using lmerTest (Kuznetsova et al., 2017) using the Satterthwaite correction for degrees of freedom. We included random intercepts and slopes for all within-person predictors to account for inter-individual variance in repeated measurements. We considered α ≤ .05 as significant. Data was visualized using ggplot2 (Hadley, 2016), ggridges (Wilke, 2017), ggpointdensity (Kremer, 2019), and ggside (Landis, 2021).

## Results

To sample stress over a period of four weeks with concurrent CGM, participants were asked to complete EMA ratings two times per day. In line with the instructions, the median lag between consecutive EMA runs was 10.85 h (interquartile range: 13.04 h). Moreover, as instructed, ratings were distributed well over different hours of the day, effectively sampling over different metabolic states (for details, see Kaduk et al., 2025).

### Everyday stress is associated with mood, but stress spikes occur independently

First, we assessed how strongly stress ratings are associated with mood since both domains are conceivably linked with metabolic states (Kaduk et al., 2025). Stress ratings showed a robust association with mood state (*r* = −.51, *b* = −.22, *t*(91) = −13.77, *p* < .001). Moreover, associations were similar for happy (*b* = −.22, *t*(83) = −7.77, *p* < .001) and sad mood (*b* = .23, *t*(91) = 7.33, *p* < .001). Run-to-run changes in EMA stress ratings were largely independent of average mood state levels (*r* = −.01, *p* = .70; LME: *b* = −.09, *t*(3561) = −.27, *p* = .79, whereas they were associated with concurrent changes in mood state (*r* = −.29; *b* = −.18, *t*(95) = −9.91, *p* < .001). This correspondence was weaker compared to average stress and average mood state ratings (*r* = −.71, *p* < .001), suggesting that average stress and average mood state ratings are more strongly associated than their fluctuations. Consequently, sudden changes in stress ratings can be more clearly differentiated from mood-related effects than average stress ratings (Figure 2).

**Figure 2.**
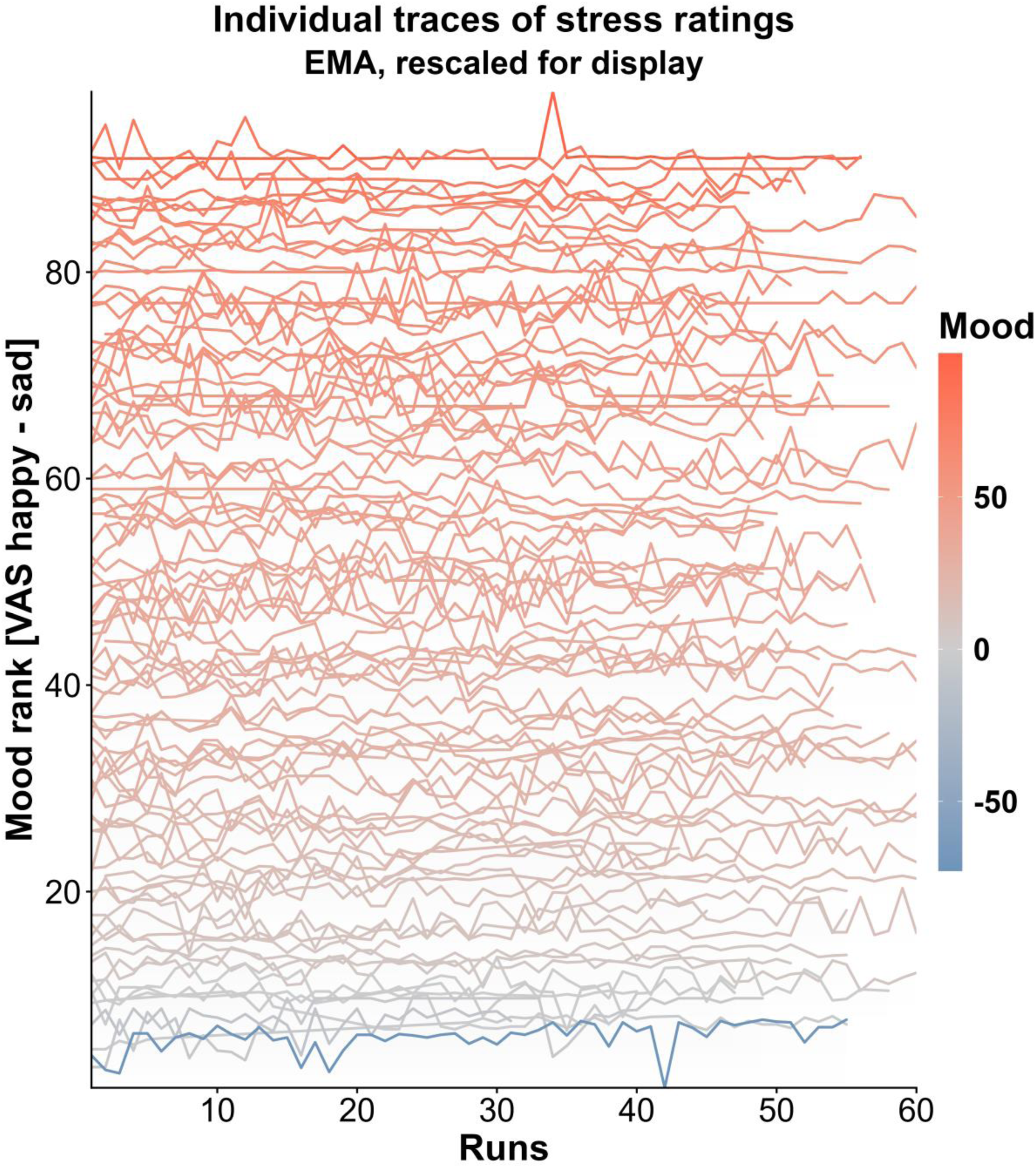
Ridgelines show individual traces of stress ratings, ranked by average mood state and color-coded accordingly. Stress ratings show a robust association with mood state ratings (*r* = −.51, *p* < .001). However, stress spikes (i.e., sudden increases and decreases; captured by the temporal derivative of stress ratings over runs) occur largely independent of typical mood state levels (*r* = −.01, *p* = .79) and were less strongly associated with sudden changes in mood state (*r* = −.29, *p* < .001). This indicates that stress spikes can be dissociated from longer-term mood-related processes.

### Higher glucose levels are associated with lower stress ratings

Next, we evaluated whether concurrent glucose levels are associated with stress ratings beyond mood. To this end, we predicted stress ratings as outcome using both group-centered and individual averages of glucose levels while controlling for positive and negative mood ratings as well as confounds. Within participants, higher glucose levels were associated with lower stress ratings (Figure 3a; *b* = −.038, *t*(711) = −2.12, *p* = .035). Individual shrinkage estimates of the effect were negative in 83 out of 91 participants (91%; Figure 3b), indicating that this association is observed in most participants. This association was not driven by individual differences in BMI or metabolic function, as indicated by weak and non-significant associations with BMI, insulin sensitivity (HOMA-IR) and the triglyceride-glucose index (*p* > .50). Sex was not associated with stress ratings (*p* = .415) and did not moderate the glucose-stress relationship (*p* = .344), suggesting that this association is consistent across male and female participants.

**Figure 3.**
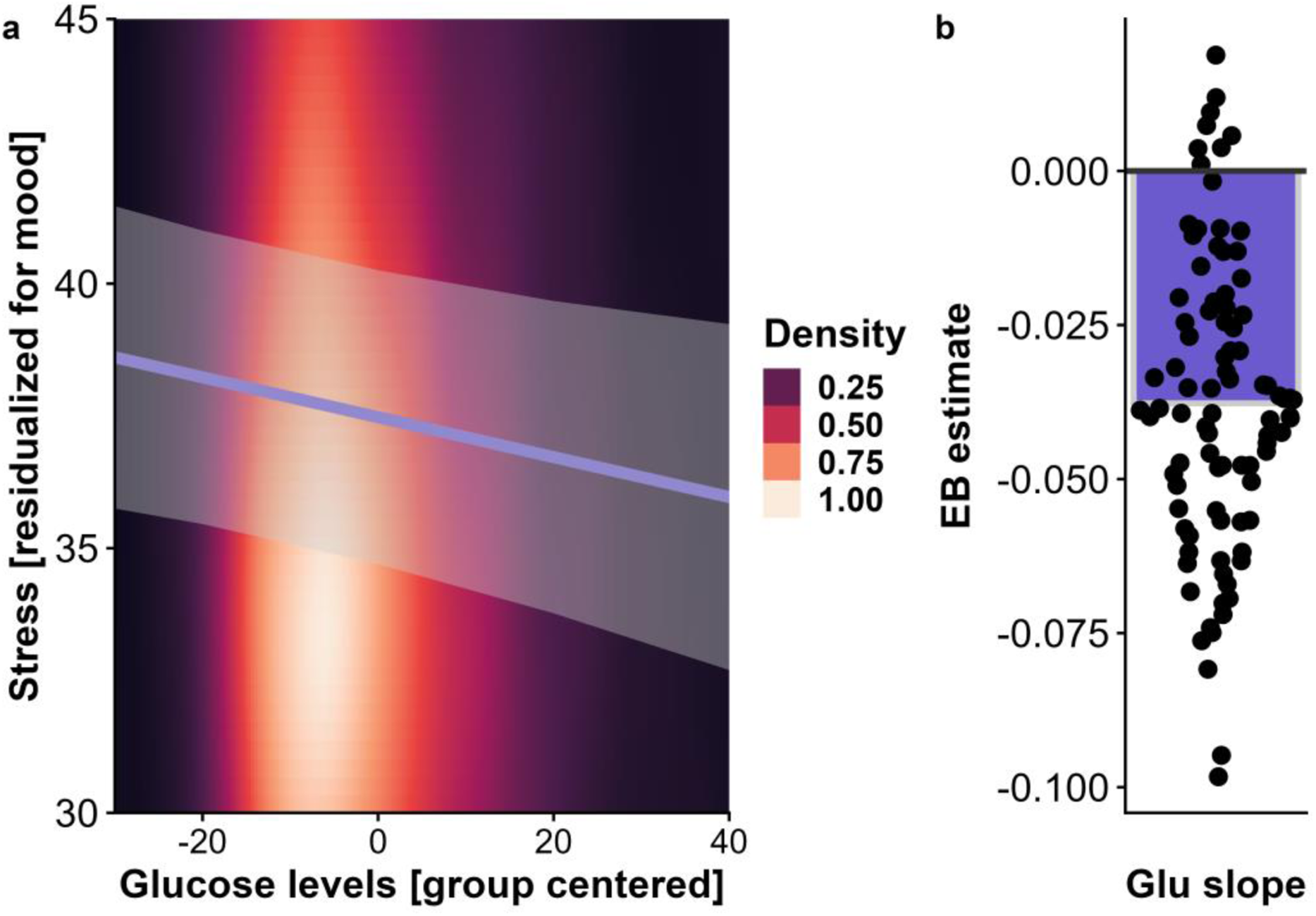
Density plot illustrating the association between glucose levels and stress ratings, controlling for mood. **A**: Stress ratings decrease as glucose levels increase. The density color scale reflects the distribution of observed values, highlighting that most data are centered near average glucose levels. **B**: Individual empirical Bayes (EB) estimates for the glucose-stress slope show that most participants exhibit a negative association, indicating lower stress at higher glucose levels (*b* = −.038, *t*(711) = −2.12, *p* = .035). Negative shrinkage estimates were observed in 91% of participants, suggesting the effect is robust across individuals.

### Differences in glucose levels precipitate changes in stress reported during EMA

To examine whether changes in glucose levels precipitate alterations in reported stress, we predicted CGM glucose levels using the time relative to the EMA stress ratings (using 2 h bins up to 12 h before the rating; see SI), the momentary change in stress and their interaction. In line with our hypothesis, we observed a significant main effect of stress change (*b* = −.37, *t*(114) = −2.53, *p* = .013), indicating that lower glucose levels in the 2 h before the stress report were associated with greater increases in stress. Additionally, we found significant interactions between time and stress change at 12 h (*b* = −.42, *p* < .001) and 8 h (*b* = −.16, *p* = .030) before EMA ratings. These interactions indicate that the changes in glucose levels during early time bins provide incremental predictive information, with individuals who later reported an increase in stress compared to the last observation (i.e., stress onset) showing a lower glucose level compared to participants experiencing stress relief (i.e., reduction in stress compared to the previous assessment) or no change in stress (Figure 4a, 4c).

**Figure 4.**
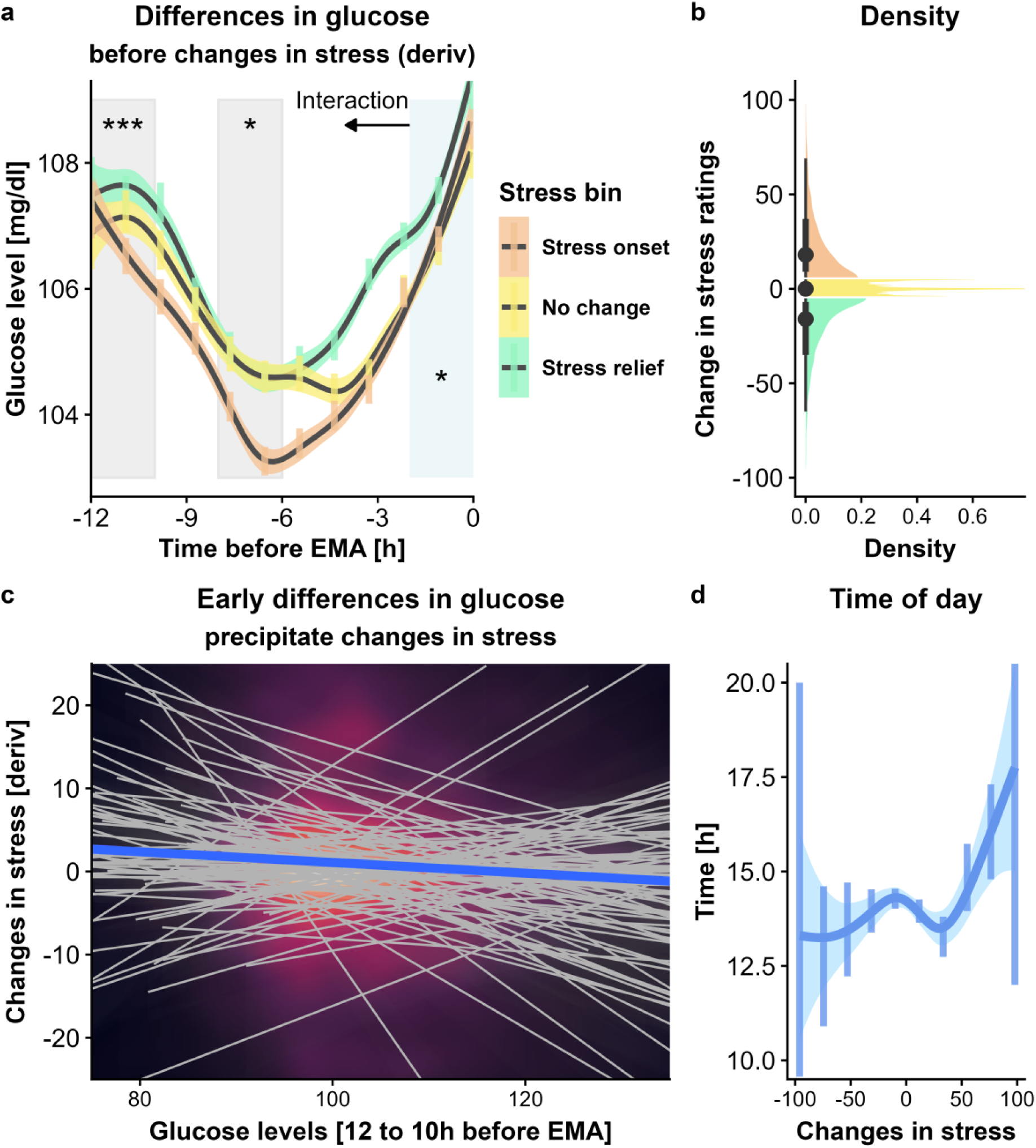
Glucose levels fluctuate before stress onset, with early drops indicating upcoming stress regardless of the time of day. **A:** Estimated glucose levels in the 12 h preceding an EMA stress rating, stratified by stress change category (stress onset, no change, stress relief). Glucose levels followed a U-shaped pattern, with drops in glucose levels starting 6-10 h before EMA. Significant interactions between time and stress change emerged at 12 h (*b* = −.42, *p* < .001) and 8 h (*b* = −.16, *p* = .030) before EMA, indicating that individuals with steeper glucose declines prior to stress reported higher subsequent stress ratings compared to those experiencing stress relief. **B**: Density distribution of changes in stress ratings across all EMA assessments. Colors indicate how the data was binned for display. **C**: Individual participant trajectories show the relationship between early glucose levels (averaged from 12 to 10 h before EMA) and subsequent changes in stress. Most participants showed that lower early glucose levels preceded greater increases in stress. **D**: Time of day distribution (24-hour format) for EMA ratings across changes in stress. Stress changes did not significantly vary by time of day (*b* = .03, *t*(159) = .32, *p* = .748). After controlling for time of day, the main effect of stress changes (*b* = −.29, *t*(121) = −2.54, *p* = .013) and the 12h bin interaction (*b* = −.26, *p* < .001) remained significant.

Since diurnal glucose variations may confound associations between glucose, stress, and affective states (La Fleur, 2003; Zink et al., 2020), we also controlled for the time of day at which EMA stress ratings were collected. We found no linear association between stress changes and time of day (*b* =.03, *t*(159) = .32, *p* = .748; Figure 4d), indicating that stress changes occurred independently of circadian patterns. To account for diurnal glucose fluctuations, we included time of day as a covariate. Even after controlling for this potential time-related variance, the main effect of stress change remained significant (*b* = −.29, *t*(121) = −2.54, *p* = .013), and the 12h bin interaction persisted (*b* = −.26, *p* < .001), though the 8h bin interaction was no longer significant (*b* = −.02, *p* = .753).

### BMI is associated with differences in stress dynamics over time

Finally, we reasoned that individual differences in energy metabolism might be associated with the dynamic changes in stress. Consequently, we next assessed whether changes in stress ratings differ according to BMI. To this end, we compared run-to-run changes in participants with normal weight, overweight, or obesity (Figure 5). Pairwise Monte-Carlo two-sample Kolmogorov-Smirnov tests revealed significant differences in the distribution of stress changes between individuals with normal weight vs. overweight (*D* = .055, *p* = .007), as well as between individuals with normal weight vs. obesity (*D* = .065, *p* = .042). In contrast, the distribution of stress changes did not differ significantly between individuals with overweight vs. obesity (*D* =.047, *p* = .302). These findings indicate that the temporal dynamics of stress changes are altered with a higher BMI, with both the overweight and obese groups showing different distributions of stress changes (i.e., more runs without any change and heavier tails of extreme changes) compared to the normal weight group. This pattern suggests that higher BMI is associated with stress ratings that fluctuate less frequently overall yet exhibit somewhat larger deviations when changes do occur.

**Figure 5.**
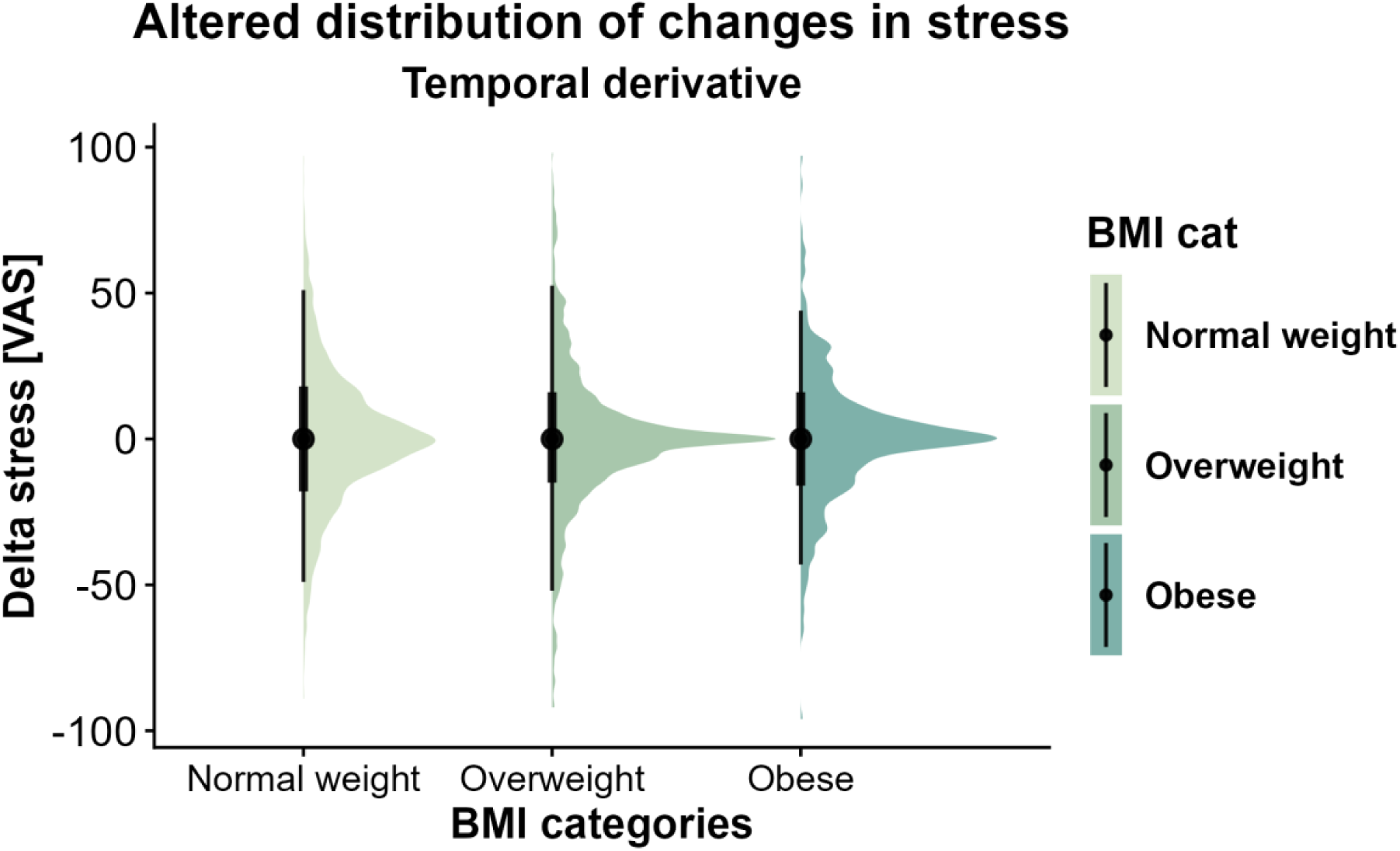
Distributions of stress changes vary with BMI. Violin plots show how moment-to-moment changes in stress (measured via VAS) are distributed for participants with normal weight, overweight, and obesity. Compared with normal weight, participants with overweight and obesity show different patterns of stress changes (normal vs. overweight: *D* = .055, *p* = .007; normal vs. obese: *D* = .065, *p* = .042). Notably, groups with a higher BMI show more pronounced peaks of minimal changes, but also heavier tails. However, there is no significant difference between the overweight and obese groups (*D* = .047, *p* = .302).

## Discussion

Adequate responses to a stressor require energy, and recent research suggests that glucose availability plays an important role in mobilizing resources to mount a sufficient response. However, it remains unclear whether naturally occurring fluctuations in glucose shape how stress is perceived in daily life by metabolically and mentally healthy individuals. By combining CGM with EMA, we found that higher glucose levels were associated with lower self-reported stress. These differences could be due to changes in allostatic behavior (i.e., eating) since stress exposure is associated with acute and subacute changes in eating that may play a functional role in stress recovery (“comfort food”). Crucially, glucose levels even preceded subsequent changes in stress reports, with lower glucose levels observed several hours before self-reported stress increases independent of time of day, suggesting that glucose patterns may serve as early markers of upcoming stress responses. Although stress and mood are closely related, acute stress spikes were only moderately associated with concurrent mood changes, suggesting that short-term stress fluctuations are sufficiently independent of mood to enable their dissociation. Overall, our findings highlight that stress is reflected not only in psychological but also in basic metabolic processes, demonstrating the large potential of combining CGM and EMA to assess stress in naturalistic settings.

Consistent with prior research, we observed a negative association between stress and mood in daily life (Richardson, 2017; Xie et al., 2024). Individuals who, on average, reported higher stress also tended to experience lower mood overall, reflected in both reduced positive and elevated negative mood. However, short-term fluctuations in stress were only weakly to moderately associated with concurrent changes in mood, suggesting that acute stress spikes may occur independently of shifts in mood (Richardson, 2017; Xie et al., 2024). This distinction between the relationship of stress and mood in general and the moment-to-moment fluctuations points to a more dynamic and context-dependent stress process at the within-person level, likely shaped by individual differences in emotion regulation (Richardson, 2017) and situational factors in daily life, such as being at work or doing physical activity (de Vries et al., 2021; Yao et al., 2024). Our findings add to recent reviews that highlight the limited research on daily and within-day stress variability, especially for non-clinical populations, despite growing evidence that stress is highly variable across time and contexts (Mengelkoch et al., 2023; Zawadzki et al., 2022). This illustrates the value and potential of combining biosensors with EMA for capturing both the stable, trait-like association between stress and mood, as well as the more transient, situational aspects of stress reactivity in daily life, including the availability of glucose as a proxy of energy levels.

Crucially, we show that stress responses are linked to glucose levels and that differences in glucose traces precipitate subsequent changes in stress for more than 12h. As expected, based on our recent review (Kördel et al., 2025), higher glucose levels were consistently associated with lower perceived stress among most participants. These results extend earlier laboratory studies showing that glucose availability modulates physiological stress responses, such as HPA-axis activity (Gonzalez-Bono et al., 2002; Kirschbaum et al., 1997; von Dawans et al., 2021), to real-world contexts and the impact on the subjective stress response. Accordingly, experimental work has demonstrated that acute glucose intake and higher glucose levels are linked with reduced tension (Benton & Owens, 1993), suggesting that glucose availability may buffer against negative affective states. We now fundamentally extend these findings by showing that the associations of glucose and stress were dynamic and preceded reported changes in subjective stress over long periods of time (up to 12h). Early drops in glucose levels precipitated the upcoming increase in reported stress, whereas elevated glucose levels were associated with subsequent reports of stress relief. These effects were observable many hours before stress reports, suggesting that glucose fluctuations may not only coincide with the momentary experience of stress but may also reflect or contribute to its regulation. This interpretation is supported by work in mice identifying fast neural pathways by which threat-related brain regions directly modulate hepatic glucose production, independent of stress hormones, suggesting that stress-related glucose dynamics are an important component of the stress response system rather than a mere by-product (Carty et al., 2025).

One possible explanation for the association of glucose traces and the subjective experience of stress involves behavioral adaptations to stress. In particular, stress can lead to changes in eating behavior (Hill et al., 2022) and has been shown to increase food cravings (Reichenberger et al., 2021). For some individuals, consuming sweet foods is associated with a reduction in psychological and physiological stress (Ulrich-Lai, 2016), potentially acting as a compensatory mechanism that affects glucose levels. However, these behavioral adaptations are not uniform since some individuals respond to stress by reducing their consumption (Hill et al., 2022; Oliver & Wardle, 1999; Torres & Nowson, 2007; Yau & Potenza, 2013). Such interindividual variability appears to be influenced by BMI (Kivimäki et al., 2006) and cortisol stress responses (Newman et al., 2007). Consistent with this idea, we observed that individuals with overweight or obesity reported more stable stress levels yet with larger extreme fluctuations (i.e., heavy tails) over time, compared to individuals with normal weight. This suggests that BMI and associated differences in energy metabolism may alter HPA axis function, leading to changes in stress experiences and dynamic adjustments (Kühnel et al., 2023). Prior work further indicates that stress, rather than general negative affect, is the more reliable predictor of eating-related responses (Reichenberger et al., 2021), proposing that stress-related eating may depend not only on momentary stress levels but also on BMI-related differences in stress dynamics. Therefore, glucose availability may not only shape stress perception through its central role in supplying energy and via HPA axis modulation, but also indirectly through behavioral changes, including stress-induced eating. Although the observed associations between glucose and stress are promising, our sampling design does not allow us to disentangle the behavioral pathways that should be addressed in future work. Taken together, our findings indicate that glucose traces effectively track changes in physiological signals that either shape or reflect emerging and ongoing changes that are linked to the onset or relief of stress in everyday life.

While this study uses an innovative combination of CGM and EMA to demonstrate that glucose levels affect self-reported stress experiences in everyday life, some limitations should be considered. First, our participants did not report whether they faced a specific stressor or stressful situation, or identify whether it was physiological, psychosocial, or psychological. Previous research suggests that the type and intensity of stressors likely modulate HPA axis responses (Kogler et al., 2015; Rüttgens & Wolf, 2022), and environmental context may further modulate daily stress perception (Yao et al., 2024). Second, our participants were not asked to provide the exact timing of when stress events occurred. This reduces our ability to establish time-order precedence between glucose fluctuations and perceived stress, limiting causal interpretation of directionality. Therefore, future studies should aim to assess stressor categories and contexts in more detail, while also collecting data regarding the onset and duration of stress experiences to better evaluate how and when glucose fluctuations relate to stress responsiveness. Accounting for individual differences in compensatory behavior like stress-eating tendencies could further clarify the mechanisms through which metabolic states shape experiences of stress. Given that glucose mobilization supports physiological arousal and emotional reactivity (Blake et al., 2001), future research should investigate whether arousal serves as a mechanism linking metabolic state to stress perception. Incorporating additional physiological measures of stress, such as heart rate and its variability (Kim et al., 2018; Kühnel et al., 2022; Kühnel, Kroemer, et al., 2020), hair cortisol (Russell et al., 2012), or continuous cortisol sampling (Kusov et al., 2023), to provide a multidimensional assessment of stress would be valuable as well. Such approaches may improve our understanding of stress vulnerability and help identify individuals at greater risk for stress-related disorders.

Energy availability plays a pivotal role in the adaptive response to stress. In this innovative study combining CGM and EMA, we show that glucose levels affect stress reports in everyday life. Crucially, these associations extended beyond momentary glucose levels, showing incremental associations with glucose levels up to 12h before changes in stress were reported. This observation highlights a unique strength of our design, suggesting that differences in glucose traces precipitate changes in stress. Therefore, our study can be seen as a starting point in considering metabolic aspects as an essential component in investigating naturalistic stress experiences, as stress experiences are intertwined with metabolic states and metabolic health. To conclude, CGM with concurrent EMA provides a promising combination of tools to study stress-related dynamics in association with bodily signals in a real-world setting. Understanding stress physiology in daily life could lead to novel stress management strategies in the long run, pointing to interventions that may improve stress reactivity by modulating metabolic signals.

## Acknowledgement

We thank Antonia Schlaich, Johanna Voß, Franziska Peglow, Hannah Groß, Sophie Mathis, David Bartsch, Rauda Fahed, Katharina Bertl, Johanna Theuer, Marie Kaeber, Ebru Sarmisak, Laura Heidiri, and Ricarda Bode for help with data acquisition and Maria Berjano Torrado for support in preprocessing the glucose data. The study was supported by the German Research Foundation (DFG) grants DE 2319/22-1, KR 4555/7-1, KR 4555/9-1; KR 4555/10-1. MK was supported by the International Research Training Group: Women’s Mental Health Across the Reproductive Years (IRTG 2804), funded by the DFG; grant number: GRK 2804/1.

## Author contributions

NBK and BD were responsible for the study concept and design. MG, KK, & AK collected data under supervision by NBK and BD. NBK conceived the method of this paper, and AK & AG preprocessed the data. NBK & MK performed the data analysis. MK & NBK wrote the manuscript. All authors contributed to the interpretation of findings, provided critical revision of the manuscript for important intellectual content, and approved the final version for publication.

